# Astrocytic PI3Kα controls synaptic plasticity and cognitive function via serine metabolism

**DOI:** 10.1101/2025.02.11.637597

**Authors:** Alba Fernández-Rodrigo, Celia García-Vilela, Abel Eraso-Pichot, Esperanza López-Merino, Irene B. Maroto, Silvia Gutiérrez-Eisman, Carla Sánchez-Castillo, Cristina Boers-Escuder, Eneko Merino-Casamayor, Mariona Graupera, María I. Cuartero, Albert Quintana, Giovanni Marsicano, José A. Esteban

## Abstract

Astrocytes are known to modulate neuronal activity by gliotransmission and through metabolic regulation. However, the connection between these two processes is still poorly defined. In this work we show that the p110α isoform of the phosphatidylinositol 3-kinase (PI3K) in astrocytes is required for long-term potentiation (LTP) and has an impact on learning and memory. Using a specific deletion of p110α from hippocampal astrocytes in adult mice, we found that LTP depends on astrocytic p110α to sustain D-serine levels for the activation of NMDA receptors during LTP induction. This requirement is based on the L-serine biosynthetic pathway of the astrocyte, which is defective in the absence of p110α because of a reduced glycolytic flux. Accordingly, the behavioral impairment in mice lacking p110α can be rescued by *in vivo* administration of L-serine. These results link for the first time the function of PI3K in astrocytes to cerebral metabolism and its influence in synaptic plasticity and cognition.

## INTRODUCTION

Neurons are able to modify their synapses upon activity they receive, altering the strength of their responses, the efficacy of synaptic transmission and, eventually, the function of neural circuits. This phenomenon is called synaptic plasticity and underlies several cognitive processes including learning and memory^1,2^. However, neurons are not the only cells contributing to synaptic plasticity. In fact, it is now well established that astrocytes are able to respond to neuronal activity and extracellular signals and, in turn, modulate neuronal and synaptic function. This occurs through the release of active molecules (gliotransmitters)^3–6^, such as glutamate^7–13^ or ATP^14–16^, directly acting on neuronal receptors. In addition, it is becoming increasingly accepted that astrocytes can also modulate neuronal function and brain activity via their role in metabolic support^17–19^. In fact, the exchange of metabolites between neurons and astrocytes is being revealed as an important contributor to neuronal function. This is best exemplified with the astrocyte-neuron lactate shuttle (ANLS)^20,21^ or with the supply of L- and D-serine from the astrocyte for the activation of neuronal NMDA receptors (NMDARs)^22–24^. These astrocyte-neuron metabolic interactions have been shown to have functional consequences for cognitive performance^25,26^ and are probably linked to human disease^27,28^. However, the intracellular mechanisms controlling this crosstalk between metabolic and synaptic regulation remain poorly understood.

In order to fill this gap, we have turned our attention to the phosphatidylinositol 3-kinase (PI3K) signaling pathway. The PI3K/mTOR pathway is well-known to mediate synaptic plasticity mechanisms^29–32^. At the same time, this is a critical hub for the regulation of cellular metabolism^33,34^, particularly controlling glucose homeostasis^35–38^ via intracellular trafficking of glucose transporters^39,40^ and by modulating the activity of glycolytic enzymes^41,42^. Therefore, the PI3K pathway appears as an attractive candidate to link metabolic regulation and synaptic plasticity. However, PI3Ks are a diverse family of signaling enzymes. Class IA PI3Ks are heterodimers composed of a p85 regulatory subunit and a p110 catalytic one, which catalyzes the production of phosphatidylinositol (3,4,5)-trisphosphate (PIP_3_)^43^. Different paralogs of the p110 catalytic subunit (commonly referred to as isoforms: p110α, p110β or p110δ) are encoded by different genes, and have been shown to play different roles in cellular physiology and growth^34,44,45^. Also, our group has recently described that neuronal p110α and p110β have separate functions in synaptic plasticity and neuronal architecture^32^. However, there is little information on regulatory mechanisms of metabolism in astrocytes, and certainly nothing known about the potential role of astrocytic PI3K signaling in these cells and its impact on neuronal function.

In order to address this question, we have used a conditional and brain region-restricted floxed mouse model for the p110α catalytic isoform of PI3K, in combination with behavioral, electrophysiological, imaging and bioenergetic assays. In light of these experiments, we now report that the lack of astrocytic p110α impairs NMDAR-dependent LTP and cognitive function due to a decrease in the availability of D- and L-serine, which can be rescued *in vitro* and *in vivo*. Moreover, we show that L-serine deficiency is caused by a decreased glycolytic flux, which is compensated by an increased mitochondrial respiration in astrocytes.

## MATERIALS AND METHODS

### Animals and ethics statement

All biosafety procedures and animal experimental protocols were approved by the Ethical Committee from the Consejo Superior de Investigaciones Científicas (CSIC), in strict accordance with Spanish (RD 53/2013, 32/2007) and EU guidelines set out in the European Community Council Directives (2010/63/EU, 86/609/EEC; PROEX 160.1/21). Animal’s health and welfare were monitored by a designated veterinarian. All mice used were housed in standard cages (maximum of 5 animals per cage) in humidity and temperature-controlled rooms. Autoventilated racks contained individually ventilated cages with absolute filters. Irradiated standard safe diet and autoclaved water were available *ad libitum*.

p110α^flox/flox^ is a transgenic mouse line with the 18^th^ and 19^th^ exons of the *PIK3CA* gene flanked by two LoxP sequences. These mice were generously donated by Bart Vanhaesebroeck (University College London, UK). Transgenesis procedures to generate these mice were previously described^46^. Their genetic background is C57BL/6J. Genotyping was carried out by Polymerase Chain Reaction (PCR) using as primers 5’-GGATGCGGTCTTTATTGTC and 5’-TGGCATGCTGCCGAATTG. For some approaches, always specified in the text, C57BL/6J-OlaHsd wild type mice were used as controls.

### Stereotaxic *in vivo* microinjections of AAVs and Sindbis viruses

AAV5-GFAP-Cre-GFP virus (serotype 5, 4.1×10^12^ virus molecules/ml, Gene Therapy Vector Core at the University of North Carolina) was delivered to the hippocampus of 2-3 months old mice through two bilateral injections (600 nl each) in the dorsal (in mm from bregma, anteroposterior (AP): −1.8, mediolateral (ML): ±1.3, dorsoventral (DV): −1.7) and ventral (AP: −2.8, ML: ±2.4, DV: −2) regions. Animals were placed in the stereotaxic frame (Harvard Apparatus) over a heating pad and immobilized using blunt ear holders. Anesthesia by inhalation of isoflurane was initiated at 2.5-3% in 2% oxygen and maintained at 1.5% during the rest of the procedure. Before the start of the procedure, the animals received analgesia by subcutaneous injection of Meloxicam (5 mg/kg dose) and ocular protection with Lipolac ophthalmic gel 2 mg/g (Bausch & Lomb, #764118). Viral particles were infused using a Hamilton Neuros syringe (Hamilton, #65460-02) inserted through small holes drilled into the skull and coupled to a micropump (Quintessential stereotaxic injector, Stoelting, #53311) at an infusion rate of 120 nl/min. After the infusion, the syringe was kept in place for 5 minutes and slowly withdrawn. The virus was allowed to express for 1 month and the accuracy of the infection was confirmed by immunofluorescence using the expression of GFP as the reporter. For neuron morphology experiments, injected mice were subjected to a second round of *in vivo* injections using the previous parameters to inject Sindbis viruses expressing GFP and animals sacrificed 24 hours later.

For mitochondrial isolation of genetically modified astrocytes, the AAV-Cre was coinfected with an AAV expressing a Cre-inducible mitoTag (serotype 1, 3.5×10^12^ viral molecules/ml, 500 nl per injection). As previously described^47^, this AAV-mitoTag expresses a *Tomm20* gene fused to three HA sequences that allows the immunoisolation of mitochondria from Cre-expressing cells.

### L-serine treatment

According to previous publications, L-serine can be safely administered with the diet through the food or the drinking water^27,48,49^. For the serine-treated animals in this study, a concentration of 5 g/L of L-serine was dissolved in the drinking water and the treatment refreshed every 2 days during three weeks. A group of untreated control animals were kept in parallel.

### Behavioral assays

Behavioral tests were conducted with 3-4 months old mice of both sexes. After handling the animals for a week, tests were performed on consecutive days. Test cages and objects were cleaned thoroughly with 70% ethanol between subjects and phases to eliminate any olfactory cues. For all behavioral assays, animals were recorded from a camera located above the testing arena and analysis was performed manually using the AnyMaze software (Stoelting).

#### Open field test

This test was used to evaluate general locomotor activity and as a habituation phase to novel object location and recognition tests. Mice were placed individually in the center of an acrylic box (40×40×40 cm) with white opaque walls at ground level. The animals were allowed to explore the arena for 15 minutes while being recorded from above the test field.

#### Y maze test

The Y maze test was performed to quantify spontaneous alternation and assess deficits or changes in spatial working memory. The apparatus consists of a Y-shaped maze with three identical enclosed arms. To perform the test, we placed each mouse in the center of the maze and recorded a single trial of 5 minutes from above. The sequence of arm entrances was quantified manually and an alternation is considered when the animal enters consecutively in the three arms. The spontaneous alternation percentage was calculated as: % spontaneous alternation = (number of alternations) x 100 / (total number of arm entrances – 2).

#### Novel object location (NOL) and recognition (NOR) tests

These tests were performed 24 hours after the open field test, which was used as the first habituation phase. For the next phases, the same open field arena was used, but in the case of the NOL, the four walls were different in order to give the mice spatial cues. In the familiarization phase of the NOR, two identical objects in terms of color, texture and shape were placed on two adjacent corners of the arena. Mice were allowed to explore the objects for 7 minutes while being recorded and, after 30 minutes one of the objects was replaced for another one with different color and texture but similar dimensions. Mice were allowed to explore the objects for 7 minutes. The objects used were a pair of drinking bottles and a pair of cylindrical glass bottles. Both groups of objects were randomly assigned as familiar or novel among the subjects of study to ensure that they showed no preference for any of them. For the NOL, the objects were a pair of culture flasks identical in shape, dimensions and texture. During the familiarization phase, both objects were placed in two adjacent corners of the arena and after 30 minutes the position of one of them was changed so that both objects were placed on the diagonal axis. Exploration of the objects was defined as close contact sniffing (with the head directed towards the object). We then calculated a discrimination index (DI) that is the time spent exploring the relocated or the novel object minus the time exploring the familiar object or the object in the familiar position divided by the total exploration time.

#### Social interaction and social memory tests

This task is used to evaluate mouse preference to establish social contact and their social interaction skills and they also rely on the mice’s innate preference for novelty. For this test we used a three chamber acrylic box with clearly divided walls with a hole to give the testing mouse access to each chamber. The test consisted of three phases. On the first phase wire cages were placed on the two outermost chambers and the testing mice were allowed to explore the arena and the cages for 5 minutes. This cages allowed visual, auditory, olfactory and minimal touch interaction. The second phase corresponds to the social interaction task and it was performed 24 h later. In this test, an unfamiliar mouse was placed under the wire cage of one of the chambers (gender and age-paired) and an inanimate object was placed on the opposite cage in an identical configuration. The unfamiliar mice had been habituated to the cages for several days in time intervals of 15 min. The subject mouse was placed in the center chamber and allowed to freely explore all three chambers for 7 minutes. The test mouse was removed and the third phase was immediately started after cleaning the arena and the wire cages thoroughly with ethanol 70% to eliminate any olfactory cues. The third phase corresponds to the social memory task, in which the inanimate object was replaced by another unfamiliar mouse and the test was performed identically as in the second phase. Behavior was recorded from a camera located above the test box and analysis was performed manually in time segments of 1 minute. We measured number of visits to each chamber, time spent on each chamber and exploration. Exploration of the targets was defined as close contact sniffing of the wire cage (with the head directed towards the cage). Sitting or standing time on the wire cages was not quantified as exploration. We also calculated the discrimination index as the time exploring the subject minus the time exploring the object divided by the total exploration time, or the time exploring the new subject minus the time exploring the familiar one divided by the total exploration time.

### Reagents

2-chloroadenosine, picrotoxin and DL-APV were purchased from Sigma-Aldrich. D-serine and L-serine were purchased from Thermo Scientific. The antibody against p110α was generated in Mariona Graupera’s laboratory. Other primary antibodies were p110β (Abcam, #ab151549), NeuN (Abcam, #ab177487), TOMM20 (Abcam, #ab186735), Akt (Cell Signaling Technologies, #2920), phospho-Akt [T308] (Cell Signaling Technologies, #2965), phospho-Akt [S473] (Cell Signaling Technologies, #4060), phospho-GSK3α/β [S21/S9] (Cell Signaling Technologies, #9331), α-Tubulin (Sigma, #T6199), GFAP (Sigma, #G3893), PHGDH (Proteintech, #14719-1-AP), GFP (ThermoFisher Scientific, #A-6455), mCherry (GeneTex, #GTX59788), GSK3α/β (Invitrogen, #44-610) and hemagglutinin-A (HA, Covance/Biolegend, #MMS-101P).

### Primary astrocytic cultures

Primary cultures of hippocampal astrocytes were obtained from 1-3 days-old p110α^flox/flox^ mice. Pups were decapitated, brains dissected and tissue mechanically disintegrated. Then, the tissue was enzymatically digested with accutase for 5 min at 37°C in DMEM F12 (Gibco). Cells were centrifuged and plated in P100 plates in DMEM F12 medium supplemented with 10% FBS, sodium pyruvate, ultraglutamine, non-essential amino acids, penicillin and streptomycin. Cells were maintained at 37°C and 5% CO_2_.

### Organotypic hippocampal slice cultures

Hippocampal slices for organotypic cultures were prepared from mouse pups (post-natal days 5 to 7) according to previously described procedures^50,51^. Mice were anesthetized and quickly decapitated once they were irresponsive to tail and foot pinches. Whole brains were extracted and immersed in ice-cold dissection solution (10 mM glucose, 4 mM KCl, 26 mM NaHCO_3_, 233.7 mM sucrose, 5 mM MgCl_2_, and 1 mM CaCl_2_ with 0.001% (w/v) phenol red as a pH indicator) previously saturated with carbogen (5-10% CO_2_, 90-95% O_2_). After dissection of the hippocampi under sterile conditions, 400 µm-thick slices were prepared using a tissue slicer (Stoelting Europe, #51425). These slices were then transferred to porous nitrocellulose permeable membranes (Merck Millipore, PICM0RG50) and maintained on culture medium (0.8% (w/v) minimum essential medium powder, 20% (v/v) horse serum, 1 mM L-glutamine, 1 mM CaCl_2_, 2 mM MgSO_4_, insulin (1 mg/liter), 0.0012% (v/v) ascorbic acid, 30 mM HEPES, 13 mM D-glucose, and 5.2 mM NaHCO_3_) at 35.5°C and 5% CO_2_ *in vitro* for 10 to 15 days. The culture medium was replaced every 2-3 days.

### Acute hippocampal slices

Acute hippocampal slices were obtained from 3-4 months old mice of both sexes. Animals were anesthetized by isoflurane inhalation and quickly decapitated once they were irresponsive to tail and foot pinches. The brains were rapidly extracted and submerged in Ca^2+^-free ice-cold dissection solution (10 mM D-glucose, 4 mM KCl, 26 mM NaHCO_3_, 233.7 mM sucrose, 5 mM MgCl_2_, and 0.001% (w/v) phenol red as a pH indicator) previously saturated with carbogen (5-10% CO_2_, 90-95% O_2_). Coronal slices (300 µm) were obtained cutting the brain in the same solution with a vibratome (Leica, VT1200S) and left in carbogen-gassed aCSF (119 mM NaCl, 2.5 mM KCl, 1 mM NaH_2_PO_4_, 26 mM NaHCO_3_, 11 mM glucose, 1.2 mM MgCl_2_, 2.5 mM CaCl_2_, and osmolarity adjusted to 290 mOsm) for 1 hour at 32°C for recovery. After that time, the slices were maintained at 25°C and the recordings were performed in aCSF.

### Viral delivery to culture systems

For organotypic slices cultures, the AAV was injected into the CA1 *stratum radiatum* region using glass pipettes. The AAV delivery was performed through pressurized nitrogen pulses by a Picospritzer III unit (Parker Instrumentation; 15 ms pulse duration) and controlled by a DG2A Train/Delay Generator (Digitimer; 2-5 Hz pulse frequency). Organotypic slices were infected the day after their preparation and the AAV expression was allowed for 10 to 15 days.

For astrocytic primary cultures, 7-days *in vitro* astrocytes were seeded in P35 plates and infected with an adenovirus CMV-Cre-IRES-mCherry (1.26×10^11^ IU/ml) or an empty vector CMV-mCherry (1.69×10^11^ IU/ml) (Viral Vector Production Unit, UPV, Universitat Autònoma de Barcelona) for 5 days before performing glucose uptake or extracellular lactate measurements.

### Electrophysiology

Excitatory postsynaptic currents (EPSCs) and field excitatory postsynaptic potentials (fEPSPs) were recorded from CA1 pyramidal neurons with recording glass electrodes while stimulating the Schaffer collateral fibers to evoke synaptic responses. During the recordings, the slices were placed in an immersion chamber constantly perfused with aCSF gassed with carbogen (5-10% CO_2_, 90-95% O_2_) and its temperature closely monitored at 29°C for whole-cell recordings and at 25°C for field recording experiments. For all experiments the aCSF was supplemented with 100 µM picrotoxin and for recordings on organotypic slice cultures also with 4 µM 2-chloroadenosine. For some recordings, 100 μM DL-APV was added to the aCSF to block NMDARs. For D-Serine and L-Serine rescue experiments, 10 µM D-Serine and 50 µM L-Serine were added to the aCSF, respectively. Patch-clamp recording glass electrodes (3-6 MΩ) were composed of silver/silver chloride electrodes inserted in glass pipettes filled with internal solution (115 mM CsMeSO_3_, 20 mM CsCl, 10 mM HEPES, 2.5 mM MgCl_2_, 4 mM Na_2_-ATP, 0.4 mM Na-guanosine triphosphate, 10 mM sodium phosphocreatine, 0.6 mM EGTA, 10 mM lidocaine N-ethyl bromide, and pH adjusted to 7.25 and osmolarity to 290 ± 5 mOsm). In the case of field recordings, glass pipettes were filled with the aCSF used as extracellular solution and placed in CA1 *stratum radiatum*.

AMPAR-mediated responses were measured as the peak amplitude of the response while the membrane potential was clamped at −60 mV. NMDAR-mediated responses were recorded by clamping the cell at +40 mV and measuring the tail response at a point when AMPAR-mediated responses had fully decayed (65 ms post-stimulation). Synaptic responses were averaged over 50–70 trials. fEPSPs were recorded at different stimulation intensities (50 µs and 20-250 µA) on each slice to generate an input-output curve and stimulation intensity was adjusted to 30% (for NMDAR-LTP and PPF) or 50% (for NMDAR-LTD) of the maximum response. At least 20 minutes of stable baseline was recorded (0.067 Hz stimulation) prior to plasticity induction. NMDAR-dependent LTP was induced with a theta burst stimulation (TBS) protocol composed of 10 trains of bursts (4 pulses at 100 Hz with a 200 ms interval) and it was repeated for 4 cycles with 20 s inter-cycle interval. NMDAR-dependent LTD was induced using a low-frequency stimulation protocol (LFS) that delivers 900 pulses at 1 Hz. Paired-pulse facilitation (PPF) experiments were performed with pairs of pulses of varying inter-stimulus intervals (50, 100, 200 and 400 ms). Data acquisition was achieved using Multiclamp 700A/B amplifiers and pCLAMP 9/10 software (Molecular Devices) and data analysis was performed with custom-made Excel (Microsoft) macros and with Clampfit software (Molecular Devices).

### Immunohistochemistry and confocal microscopy

Mice were anesthetized by isoflurane inhalation and intracardially perfused with 0.1 M pH 7.4 phosphate buffer (PB) to wash the vascular system prior to injecting the fixative (4% paraformaldehyde in PB pH 7.4). Brains were post-fixed overnight immersed in the same fixative solution at 4°C and then sequentially dehydrated in solutions containing 15 and 30% sucrose in PB at 4°C until they sank. Brains were cut in 50-75 µm coronal slices in cold PBS with a vibratome and then blocked with 5% bovine serum albumin (BSA) (Sigma, #A4503) and 0.25% Triton X-100 (BioRad, #1610407) in PBS for 2 h at room temperature. Once blocked, slices were incubated with the primary antibody in blocking solution at 4°C overnight. After several washes, slices were incubated with the secondary antibody, being Alexa 488, 555 or 647 (ThermoFisher, provided by the microscope service facility) depending on the experiment. Finally, they were stained with 4′,6-diamidino-2-phenylindole (DAPI; 1 μg/ml) for 5 minutes and mounted on adherent microscope slides (Thermo Scientific, #15438060) with Prolong Glass Antifade (ThermoFisher, #P-36982). Fluorescence images were acquired as z-stacks with a confocal inverted microscope (LSM800, Zeiss) using a 20x NA 0.8 Plan-Apochromat or 40x NA 1.3 oil Plan-Apochromat objectives and 488-nm, 555-nm and 647-nm lasers in combination with ZenBlue 3.3 software. Z-stacks were reconstructed (maximum intensity projection) using Fiji v1.54f. For animals infected with AAV5-GFAP-Cre-GFP, percentage of infection was quantified as the ratio of GFP-positive cells over the total GFAP-positive astrocytes. For neuronal morphology studies, neuronal dendritic trees were two-dimensionally traced with the simple neurite tracer plug-in. Sholl analysis was performed tracing the number of intersections in 10 µm intervals from the soma, and the total basal and apical dendrite length was also measured. For spine morphology, images were acquired using a 63x NA 0.8 Plan-Apochromat M27 oil immersion objective and a 488 nm laser. Spine density was defined as the number of spines divided by the dendritic length.

### Electron microscopy

Mice were anesthetized by isoflurane inhalation and intracardially perfused with PB prior to injecting the fixative (4% paraformaldehyde, 2% glutaraldehyde in 0.1 M PB pH 7.4). Brains were left for 2 h at room temperature (RT) and overnight at 4°C immersed in the same fixative solution. Coronal slices of 200 µm were obtained with a vibratome and post-fixed in a 2% solution of osmium tetroxide in 0.1 M PB (1.5 h, RT). The slices were then washed and stained with 2% uranyl acetate in water (1 h, RT in darkness), washed, dehydrated in increasing concentrations of ethanol and embedded in Epon resin. Series of ultrathin sections of the CA1 and *stratum radiatum* (SR) regions of the hippocampus were collected and mounted on single oval slot copper/palladium grids. The sections were imaged on a JEM-1400 Flash transmission electron microscope (JEOL) coupled to a CMOS Oneview camera (Gatan). The SR was imaged with a 6,000-8,000x magnification to quantify mitochondrial density and a 20,000x-30,000x magnification for mitochondrial morphology, measured using Fiji v1.54f as previously described^52^. We measured the mitochondrial area, the aspect ratio (major/minor axis length) and the length and width of the cristae.

### Western blotting

Whole hippocampi were homogenized in ice-cold lysis buffer containing 150 mM NaCl, 50 mM Tris-HCl (pH 7.5), 1 mM EDTA, 1% Triton X-100, Complete EDTA-free and PhosSTOP and centrifuged at 13200 rpm for 5 min at 4°C. The resultant supernatant was processed by SDS-polyacrylamide gel electrophoresis and transferred to polyvinylidene fluoride (PVDF) membranes (Immobilon-P, Merck Millipore). Membranes were blocked for 1 h in 5% BSA in Tris-buffered saline with 0.1% Tween-20, then incubated with primary antibodies overnight at 4°C in blocking solution. Membranes were washed and incubated for 1 h at room temperature in the corresponding horseradish peroxidase-conjugated (HRP) secondary antibodies (Jackson ImmunoResearch) prepared in blocking solution. Immunodetection was carried out by chemiluminescence (Immobilon Western, Millipore, WBKLS0500) using ImageQuant™ LAS 4000 mini biomolecular imager (GE Healthcare Life Sciences). The ImageJ software was used to analyze and quantify signal intensities obtained from digital images.

### Oxygen consumption rate experiments

#### Isolation of mitochondria by immunoaffinity purification

Mitochondria were isolated from freshly collected hippocampus from AAV-mitoTag and AAV-Cre coinfected mice. Tissue homogenates were prepared using the Mitochondria Extraction Kit-Tissue (Miltenyi Biotec, #130-097-340) according to manufacturer’s instructions. Briefly, following decapitation, brains were extracted and hippocampi dissected, minced with dissection scissors and digested with the manufacturer’s extraction buffer for 30 minutes on ice. After centrifugation at 300×*g* for 5 min at 4°C, pellets were resuspended in Protease Inhibition Buffer supplemented with EDTA-free protease inhibitor cocktail (Roche, #05892791001) and dissociated with a GentleMACS tissue dissociator (Miltenyi Biotec, #130-093-235). Homogenates were centrifuged at 500×*g* for 5 min at 4°C and the supernatants containing mitochondria were purified as directed in the Mitochondria Isolation Kit, mouse tissue and the µMACS HA Isolation Kit (Miltenyi Biotec, #130-096-946 and #130-091-122). For magnetic labelling and isolation of the mitochondria, supernatants diluted in Separation Buffer were incubated with 50 μL of anti-HA Microbeads on a rotation wheel for 2 h at 4°C. Subsequently, LS columns (Miltenyi Biotec, #130-042-401) were placed in a magnetic QuadroMACS^TM^ Separator (Miltenyi Biotec, #130-090-976) and equilibrated with 2 ml Separation Buffer. Microbead-coated mitochondria were applied onto the LS column, followed by three 3-ml washing steps with Separation Buffer. Columns were removed from the separator and mitochondria gently flushed out in 1.5 ml of Separation Buffer with a plunger. Mitochondria were pelleted by centrifugation for 2 min at 13,000×g and resuspended in cold Isolation Buffer.

#### Oxygen consumption rate measurements

Oxygen consumption rate (OCR) experiments were carried out in an Oroboros oxygraph-2k (Oroboros Instruments) in the MiR05 media (0.5 mM EGTA, 3 mM MgCl_2_, 60 mM Lactobionic acid, 10 mM KH_2_PO_4_, 20 mM HEPES, 110 mM Sucrose, 1 g/L BSA, fatty acid free, pH 7.1). Freshly isolated mitochondria (30 µl) were injected in each chamber and a mix of 2 mM malate, 5 mM pyruvate and 10 mM glutamate was used as starting substrates for complex I, followed by 1.25 mM ADP+Mg^2+^. Subsequently, 10 mM succinate was added as complex II substrate. Then, different inhibitors of the mitochondrial complexes were added: 2.5 µM oligomycin (complex IV), 0.5 µM rotenone (complex I) and 2.5 µM antimycin A (complex III). The uncoupler carbonyl cyanide 3-chlorophenyl (CCCP) was used in 1 µM steps until no respiration increase was observed. In some experiments 10 µM cytochrome c was used as a mitochondrial viability control. The analysis was performed using the Oroboros DatLab software version 7.3.0.3 (Oroboros Instruments) measuring the respiration from one-minute stable regions.

### Metabolite measurements

#### Glucose uptake assay

Glucose uptake from astrocytic cultures was measured using the Glucose Uptake Assay Kit (Abcam, #ab136955). In brief, astrocytes were starved in a serum-free medium for 6 h and then kept in a glucose free Krebs-Ringer-Phosphate-Hepes buffer (20 mM HEPES, 5 mM KH2PO_4_, 1 mM MgSO_4_, 1 mM CaCl_2_, 136 mM NaCl, 4.7 mM KCl, pH 7.4) and 2% BSA. After 40 min, astrocytes were treated with10 mM 2-deoxyglucose (2-DG) for 10 min. Then, whole-cell extracts were prepared and 2-DG quantification was carried out as described by the manufacturer’s instructions.

#### Extracellular lactate

Culture medium was deproteinized with 6% perchloric acid (HClO_4_) and kept on ice for 1 h. Samples were centrifuged for 5 minutes at 4°C and the pH adjusted to 6-7 with 20% KOH. Samples were centrifuged again to remove the precipitate and the supernatant was mixed with lactate assay medium (1 M glycine, 0.4 M hydrazine-SO_4_, 1.3 mM EDTA, pH 9.5) and 7.4 mM NAD^+^ freshly prepared. The absorbance was measured in a spectrophotometer at OD 340 nm (to detect basal NADH) (E_0_) and then 10 µg/ml of lactate dehydrogenase (Roche, #10127876001) was added to convert all the lactate in the media to pyruvate with the production of NADH + H^+^. After 40 minutes of incubation at room temperature, the absorbance was read again at 340 nm (E_1_). To calculate the lactate concentration in the media we used the following equation: c= (E_1_ – E_0_)/(ε*l), being c the lactate concentration (M), ε the extinction coefficient for NADH (6.22*10^-3^ M^-1^ cm^-1^) and l the length of the cuvette (cm).

#### D-serine in hippocampal extracts

D-Serine was measured with the DL-Serine Assay Kit (Abcam, #ab241027) following the manufacturer’s instructions. Briefly, animals were anesthetized by isoflurane inhalation and quickly decapitated once they were irresponsive to tail and foot pinches. The brains were rapidly extracted and the hippocampi dissected in ice-cold aCSF previously saturated with carbogen (5-10% CO_2_, 90-95% O_2_). The hippocampi were homogenized, deproteinized and processed according to manufacturer’s instructions. D-Serine concentration is calculated from a fluorescence signal (Excitation/Emission = 535/587 nm) against a standard curve of known D-serine concentrations.

### Statistical analysis

Results were calculated as means ± standard error of the mean (SEM). Statistical differences between groups were determined via different non-parametric tests and the cut-off value for statistical significance was set at p-value<0.05. For pairwise comparisons, Mann-Whitney test was used for unpaired data and Wilcoxon test for paired data. For multiple comparisons, we used two-way ANOVA (if indicated repeated-measures), followed by Bonferroni’s post-hoc analysis; or Kruskal-Wallis, followed by Dunn’s multiple comparisons test. The values for n, p and the specific statistical test performed for each experiment are described in the corresponding figure and legends. All statistical analysis and graphic representations were performed using GraphPad Prism software v8.3.

## RESULTS

### p110α knock-out in astrocytes alters dendritic spines in hippocampal neurons

To study the role of the p110α catalytic isoform of PI3K in astrocytes we used a gene-targeted approach using Cre-lox recombination. Given the embryonic lethality of the complete knock-out of *PIK3CA* gene^53^, we performed bilateral stereotaxic hippocampal injections with an adeno-associated virus (AAV) expressing the Cre recombinase under the glial fibrillary acidic protein (GFAP) promoter (AAV5-GFAP-GFP-Cre) in the hippocampus of adult p110α^flox/flox^ mice (Fig. 1A). Saline-injected animals were used as control. With this experimental paradigm, after one month of infection, Cre was expressed in around 70-80% of the astrocytes in the *stratum radiatum* across the whole hippocampus (Suppl. Fig. 1A). The specificity of the infection for astrocytes was confirmed from the colocalization of Cre-GFP with a GFAP antibody, and not with anti-NeuN immunostaining (Fig. 1A). This approach produced a significant reduction in p110α expression levels in infected animals (referred to as p110α^aKO^, aKO stands for astrocytic knock-out) as compared to controls, without detectable changes in another catalytic subunit isoform, p110β (Suppl. Fig. 1B, C). Of important note, the knock-out efficiency was not expected to be complete, as p110α is ubiquitously expressed and is also present in neurons and cerebral vasculature^46^. This decrease in p110α levels was accompanied by a significant reduction in the phosphorylation of S473 residue of AKT and a slight but significant increase in the total AKT expression levels (Suppl. Fig. 1D, E). Importantly, astrocytic viability did not appear to be altered by p110α depletion, according to the density of GFAP-positive cells in infected animals (Fig. 1B).

**Figure 1.**
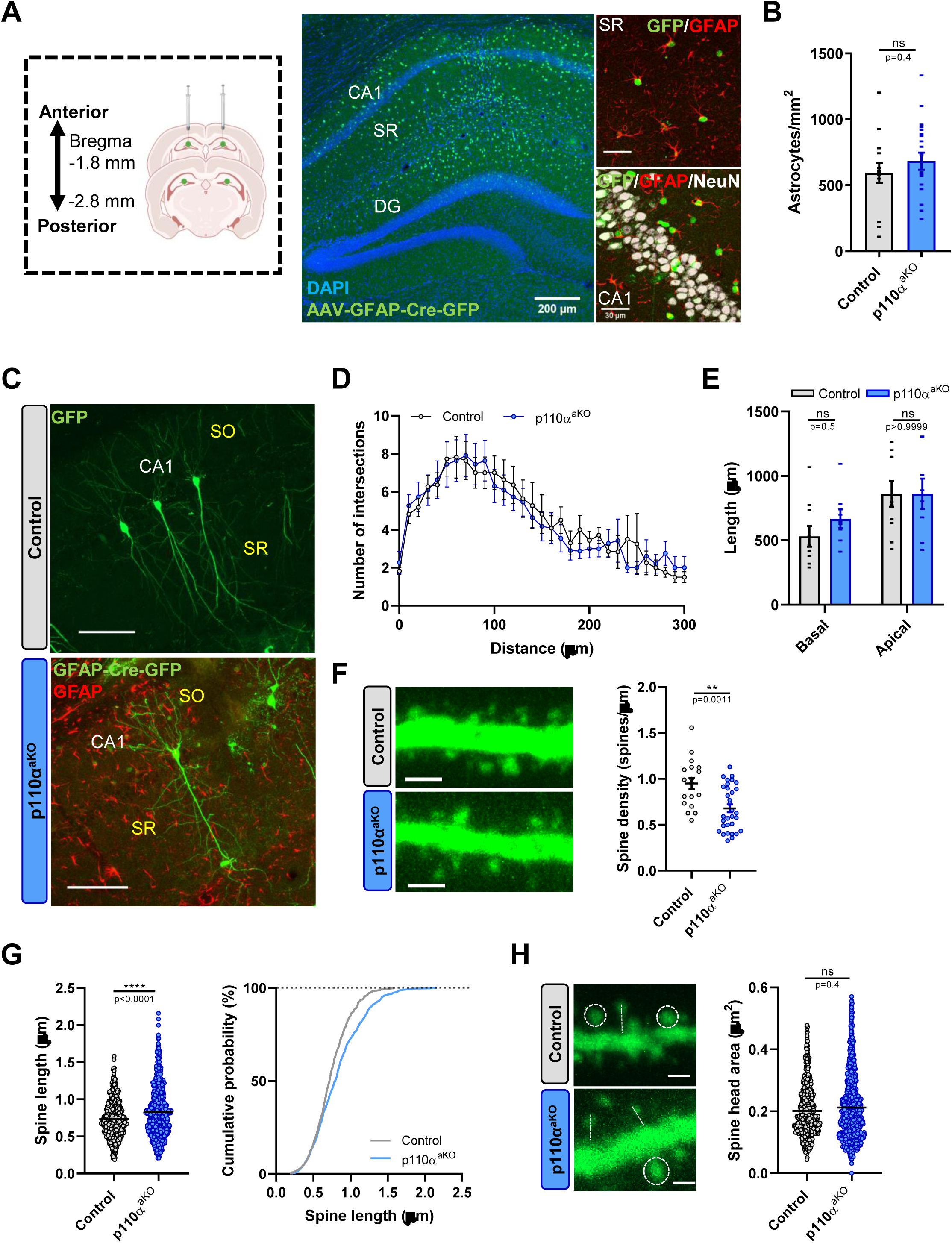
Cre-LoxP mediated conditional knock-out of p110α in the astrocytes of the hippocampus and neuronal effects. **(A)** Left, experimental scheme showing the injection sites in the hippocampus for the ablation of p110α from p110α*^flox/flox^* mice to generate the p110α^aKO^. Center, merged representative image of GFAP-Cre expression (shown in green) in the hippocampus after one month of AAV infection in combination with the nuclear marker DAPI (shown in blue). The CA1, SR (*stratum radiatum*) and DG (dentate gyrus) regions are depicted. Scale bar: 200 µm. Right, high magnification images of SR (up) and *stratum pyramidale* (down) subregions of CA1. AAV infected astrocytes are labeled by GFP (shown in green), all the astrocytes by GFAP (shown in red), the layer of pyramidal neurons in CA1 by NeuN (shown in gray) and the nuclei by DAPI (shown in blue). Scale bar: 30 µm. **(B)** Quantification of the number of GFAP^+^ astrocytes per mm^2^ in the p110α^aKO^ and control conditions. Bars represent mean±SEM. Control n=14, p110α^aKO^ n=21 slices from 4-9 mice. **(C)** Representative images of CA1 GFP-expressing neurons from p110α^flox/flox^ mice for control and p110α^aKO^ conditions. After one month of infection, mice were reinfected with a Sindbis GFP virus. Scale bars: 100 µm. **(D)** Sholl analysis depicting the mean number of intersections along the dendritic tree for control and p110α^aKO^ neurons. Statistical significance was calculated by two-way repeated-measures ANOVA. **(E)** Total dendritic length for apical and basal dendrites from GFP-infected neurons. Control n=10, p110α^aKO^ n=8 neurons from 3-4 mice. Statistical analysis was carried out with a two-way ANOVA and Bonferroni post hoc tests (ns: not significant). Data are displayed as means ± SEM. **(F)** Left, representative confocal images of GFP dendrites and spines for each condition. Scale bars: 2 µm. Right, quantification of spine density from CA1 neuron dendrites control and p110α^aKO^ mice. Control n=18, p110α^aKO^ n=32 dendrites from 4-5 mice per condition. **(G)** Left, spine length from GFP neurons and in the right their cumulative distributions of p110α^aKO^ and control mice. Control n=660, p110α^aKO^ n=1009 spines from 4-5 mice per condition. **(H)** Left, representative images of GFP spines and dendrites. Circles depict the spine area and lines the length. Right, quantification of mean spine head area from GFP neurons of p110α^aKO^ and control mice. Control n=701, p110α^aKO^ n=1070 spines from 4-5 mice per condition. (F-H) Statistical analysis was carried out by Mann-Whitney test (** p<0.01, **** p<0.0001, ns: not significant).

To study neuronal morphology in the absence of astrocytic p110α, p110α^aKO^ mice were sparsely infected with a GFP-expressing Sindbis virus for 24 h to label individual CA1 pyramidal neurons (Fig. 1C). The lack of p110α in astrocytes did not affect dendritic arborization or total dendritic length (Fig. 1D, E). However, there was a strong decrease in spine density (Fig. 1F) that was accompanied by a small but significant increase in spine length (Fig. 1G), without changes in spine head area (Fig. 1H). To note, this phenotype is clearly distinct from the one observed upon deletion of neuronal p110α, which produced a strong dendritic atrophy and an increase in spine head area^32^.

### Astrocytic deletion of p110α causes spatial and object recognition memory impairment

Previous studies from our laboratory dissected specific behavioral deficits upon deletion of neuronal p110α^32^. Therefore, we wanted to explore the role of the astrocytic p110α isoform in cognitive function and, in particular, in learning and memory. After one month of hippocampal infection, p110α^aKO^ and controls were evaluated with different behavioral tests.

First, general locomotor activity when mice are exposed to a novel environment was evaluated in the open field test (Fig. 2A). No abnormal locomotor behavior regarding speed, distance travelled or preference for different areas of the arena was observed in the p110α^aKO^ mice compared to their controls (Fig. 2A, B). To assess possible deficiencies in working memory, we studied spontaneous alternation performed in the Y maze. The spontaneous alternation percentage was statistically higher than chance for both controls and knock-out animals, without significant differences between them (Fig. 2C).

**Figure 2.**
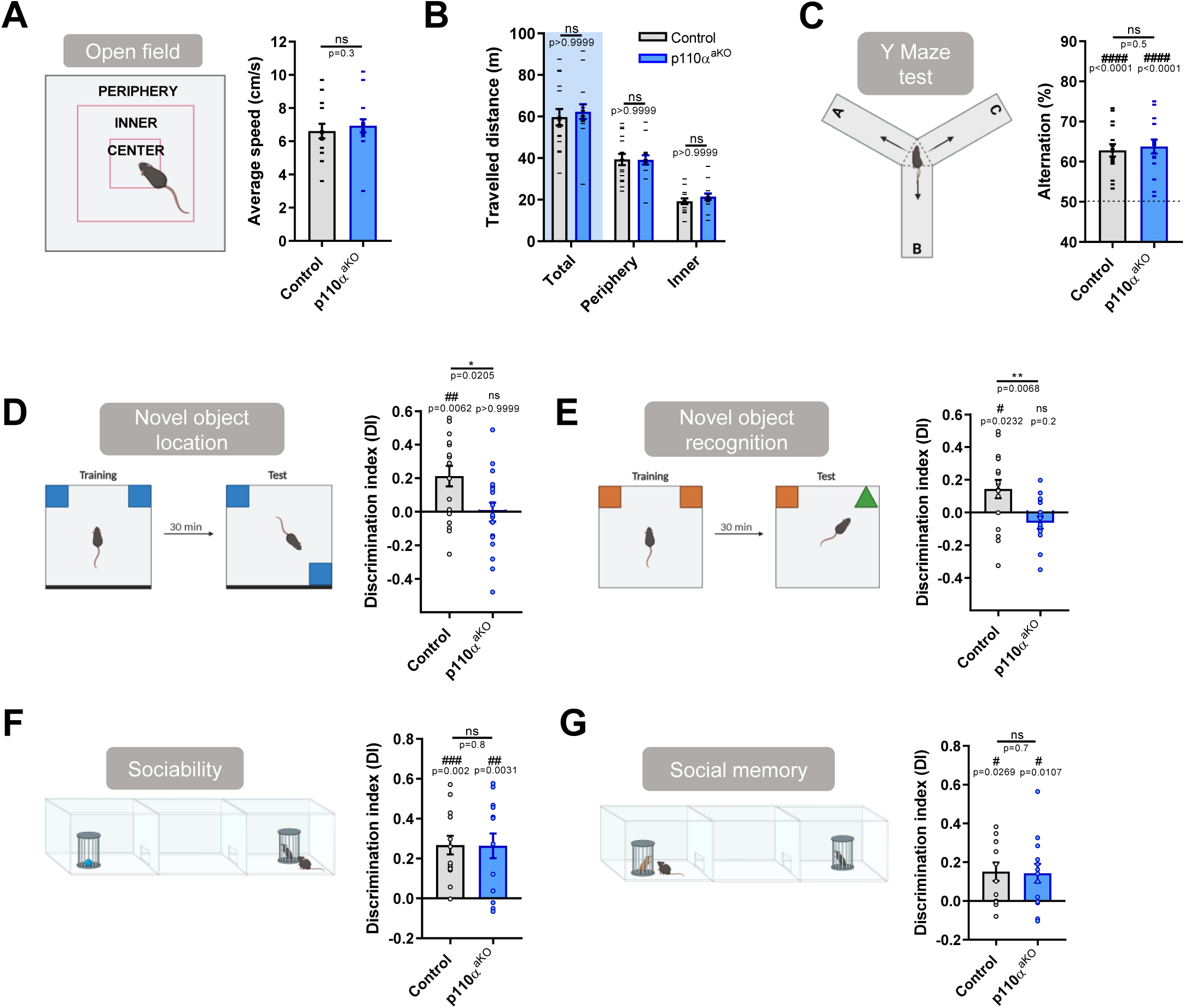
Astrocytic p110α contributes to spatial memory and object recognition. **(A, B)** Mice were tested in the open field to study locomotor behavior. Mice were placed individually in the center of an open field arena and allowed to freely explore for 15 minutes. **(A)** Left, representative picture of the open field arena and the three different areas in which it was divided; from outside to inside: periphery, inner and center. Right, average speed for the entire duration of the test for p110α (left) astrocytic knock-out mice. **(B)** Quantification of the travelled distance for the entire duration of the test separated by the peripheral and the inner+center zones of the arena for the p110α astrocytic knock-out mice. The total distance travelled is also shown (shaded background). Bars represent mean±SEM and individual values for each mouse are plotted as aligned dot-plots. Two-way ANOVA and Bonferroni post-hoc tests were used to assess statistical differences between p110α^aKO^ and their controls (ns: not significant). Control n=16, p110α^aKO^ n=16. **(C)** Left, experimental scheme of the spontaneous alternation test performed in a Y-shaped maze. Right, spontaneous alternation percentages from p110α^aKO^ and control animals. Increases on the percentage of alternation over chance were evaluated with Wilcoxon Signed Rank Test (## p<0.01, ### p<0.001). Control n=16, p110α^aKO^ n=16. **(D, E)** Left, experimental diagram of the novel object location test (D) and the novel recognition test (E). Right, discrimination indexes for the novel versus the familiar location (D) or the novel versus the familiar object (E) in p110α^aKO^ mice. Control n=17-19, p110α^aKO^ n=17-18. **(F, G)** Left, schematic representation of the three-chamber social interaction task for the sociability (F) and for the social novelty (G) tests. Right, discrimination indexes for the social target rather than the inanimate object (F) and for the novel subject rather than the known acquaintance (G) for p110α^aKO^ mice. Control n= 14, p110α^aKO^ n=14. (A, C-G) Individual values for each mouse and condition are represented along with the mean±SEM as aligned dot-plots. To assess whether the DIs were different from zero we performed the Wilcoxon Signed Rank Test (# p<0.05, ## p<0.01, ### p<0.001) and Mann-Whitney test was used to assess statistical differences between p110α^aKO^ and their controls (* p<0.05, ** p>0.01, ns: not significant). Images in (A, C-G) created with BioRender.

Next, we used the novel object location (NOL) and the novel object recognition (NOR) tests that have been widely used to study various aspects of learning and memory (see Materials and Methods). At the NOL test, while the controls were able to significantly discriminate the displaced object, p110α^aKO^ mice failed to do so (Fig. 2D). Similarly, at the NOR test, we observed that p110α^aKO^ mice were unable to discriminate between the familiar and the novel object compared to their controls (Fig. 2E). Therefore, these results indicate that p110α in hippocampal astrocytes is required for spatial memory and for object recognition memory.

Finally, we tested sociability and preference for social novelty consecutively, using a three chamber arena (see Materials and Methods). Using these tests, we observed that both controls and p110α astrocytic knock-out mice significantly preferred social exploration (Fig. 2F) and the novel subject rather than the familiar one (Fig. 2G). These results indicate that astrocytic p110α is not required for sociability or for social memory.

### The lack of p110α in astrocytes specifically impairs long-term potentiation

The behavioral deficits we have observed in p110α^aKO^ mice suggest that astrocytic p110α may also participate in synaptic plasticity paradigms related to learning and memory. To test this hypothesis, we evaluated synaptic transmission at Schaffer collateral-CA1 synapses from acute hippocampal slices using extracellular field recordings of excitatory postsynaptic potentials (fEPSPs).

Long-term potentiation (LTP) elicited by theta burst stimulation (TBS) was strongly reduced in p110α^aKO^ mice, as compared to control animals, even though it was not completely abolished (Fig. 3A). This result indicates that astrocytic p110α is required for LTP. In contrast, long-term depression (LTD) elicited by low frequency stimulation was indistinguishable between p110α^aKO^ and control mice (Fig. 3B). Furthermore, basal synaptic transmission was unaltered in p110α^aKO^ hippocampal slices compared to their corresponding controls, as assessed by input/output curves of fEPSPs at different stimulation intensities (Fig. 3C) or by calculating the ratio between AMPAR- and NMDAR-mediated EPSCs (AMPA/NMDA ratios; Fig. 3D) using organotypic hippocampal slices infected with AAV5-GFAP-GFP-Cre for 14 days.

**Figure 3.**
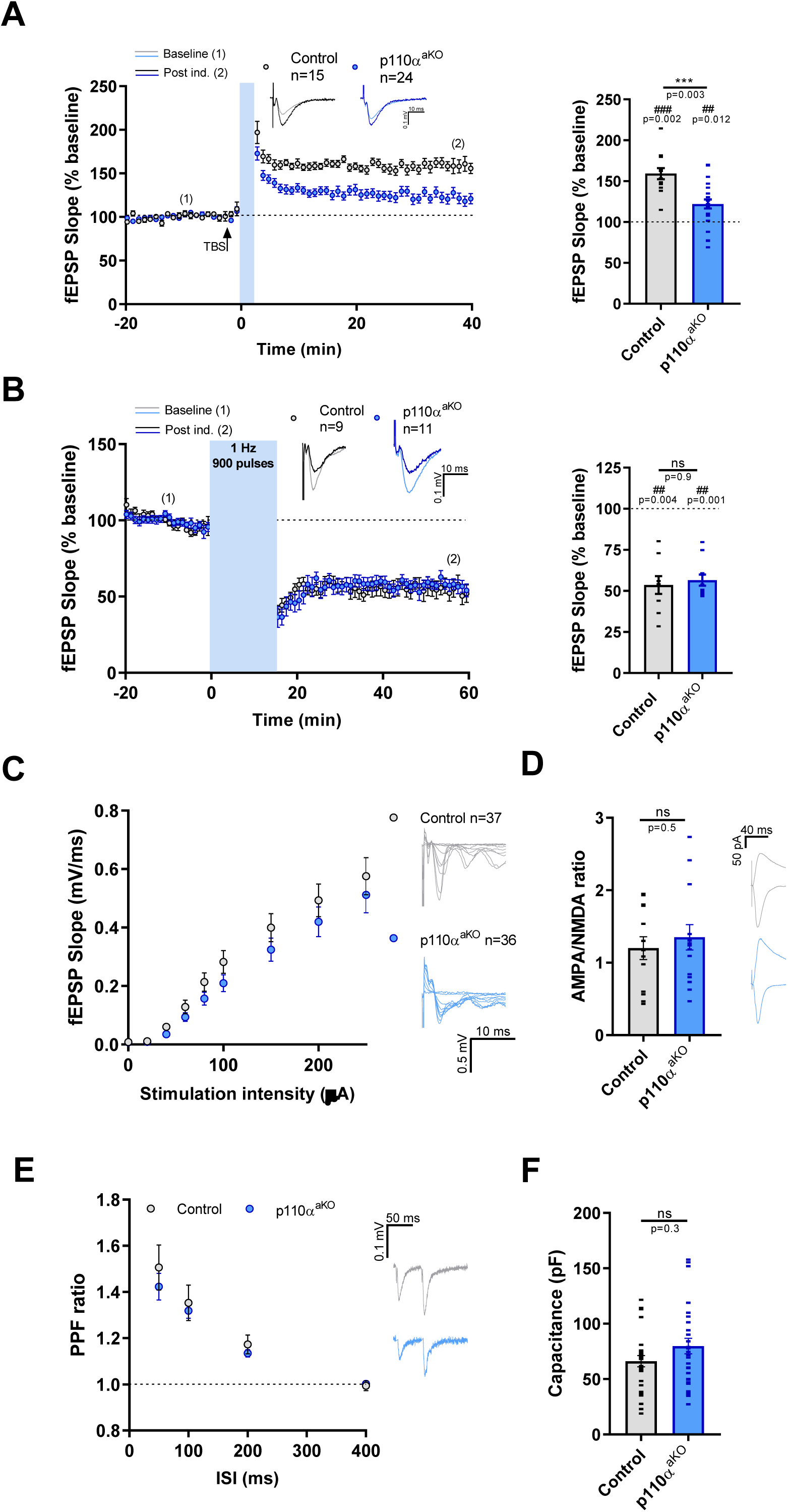
p110α^aKO^ mice have an impairment in NMDAR-LTP. **(A)** Left, time course of NMDAR-dependent LTP induced by TBS. Right, bar plot representing the mean of the fEPSP slope for each slice on the last 5 min of recording. **(B)** Left, time course of NMDAR-dependent LTD induced by LFS. Right, bar plot representing the mean of the fEPSP slope for each slice on the last 5 min of recording. (A, B) Representative traces for each condition are depicted in the upper part of the graph, the lighter color corresponding to the average trace of the baseline and the darker one to the post-induction. **(C)** Input-output curve of fEPSPs evoked at different stimulation intensities. Representative traces from one experiment for each condition are shown in the right part of the graph. Scale bar: 0.5 mV, 10 ms. **(D)** AMPA to NMDA ratio in EPSCs recorded from CA1 pyramidal neurons from p110α^aKO^ and controls. AMPA transmission is measured at −60 mV and NMDA approximately 60 ms after the onset of the stimulation at +40 mV. Control n=13, p110α^aKO^ n=15 cells. Representative traces are shown in the right. Scale bar: 50 pA, 40 ms. **(E)** Pairs of pulses separated 50, 100, 200 and 400 ms are delivered to the Schaffer collaterals for measuring the paired-pulse ratio (PPF). Control n=8, p110α^aKO^ n=16 from 7-8 mice. Representative traces for the 50 ms inter-stimulus interval are shown in the right. Scale bar: 0.1 mV, 50 ms. **(F)** Whole-cell membrane capacitance. Control n=29, p110α^aKO^ n=31 cells. Bars represent mean±SEM and individual values for each condition are plotted as aligned dot-plots. Statistical differences were assessed by two-way repeated-measures ANOVA with Bonferroni post-hoc test (C, E) and Mann-Whitney test (A, B, D, F) (p>0.001, ns: not significant). Wilcoxon Signed Rank Test was performed to assess statistically significant potentiation (A) or depression (B) (## p>0.01, ### p<0.001). n, number of slices per condition from n>6 mice per condition.

We also evaluated presynaptic function in acute hippocampal slices by measuring paired pulse facilitation. This was unaffected too at all inter-stimulus intervals (Fig. 3E), suggesting that the absence of astrocytic p110α does not significantly alter presynaptic glutamate release at CA3-to-CA1 synapses. Finally, whole-cell capacitance, as an indicator of total cell surface, was not altered in CA1 neurons from p110α^aKO^ animals (Fig. 1F).

### The LTP impairment of p110α^aKO^ mice is accompanied by a deficient NMDAR activation and can be rescued by D- and L-serine

Given the specific deficit in LTP caused by the absence of astrocytic p110α, we asked whether LTP induction may be altered in p110α^aKO^ animals. To address this point, we analyzed the fEPSP response during the 4 pulses at 100 Hz of the TBS protocol used for LTP induction. Specifically, we measured the fEPSP amplitude 60 ms after the last pulse, which would correspond mostly to the NMDAR response (blue and black arrows in Fig. 4A). Using this analysis, we observed that this slow fEPSP gradually increased in control animals across the 4 rounds of TBS of the LTP induction protocol (Fig. 4B, gray symbols). In contrast, this response was consistently smaller and did not increase in p110α^aKO^ animals (Fig. 4B, blue symbols), being significantly smaller than that of control animals at the fourth induction round (Fig. 4B, C). To note, similar experiments carried out in the presence of D, L-2-amino-5-phosphonovaleric acid (APV; NMDAR antagonist) in control slices showed a marked reduction in this slow fEPSP, indicating that it indeed corresponds mostly to NMDAR activation (Fig. 4B, yellow symbols). Given the normal synaptic transmission mediated by AMPARs and NMDARs in p110α^aKO^ animals (Fig. 3C, D), these results strongly suggest that the absence of astrocytic p110α produces a deficient NMDAR response specifically during the high demanding conditions of LTP induction.

**Figure 4.**
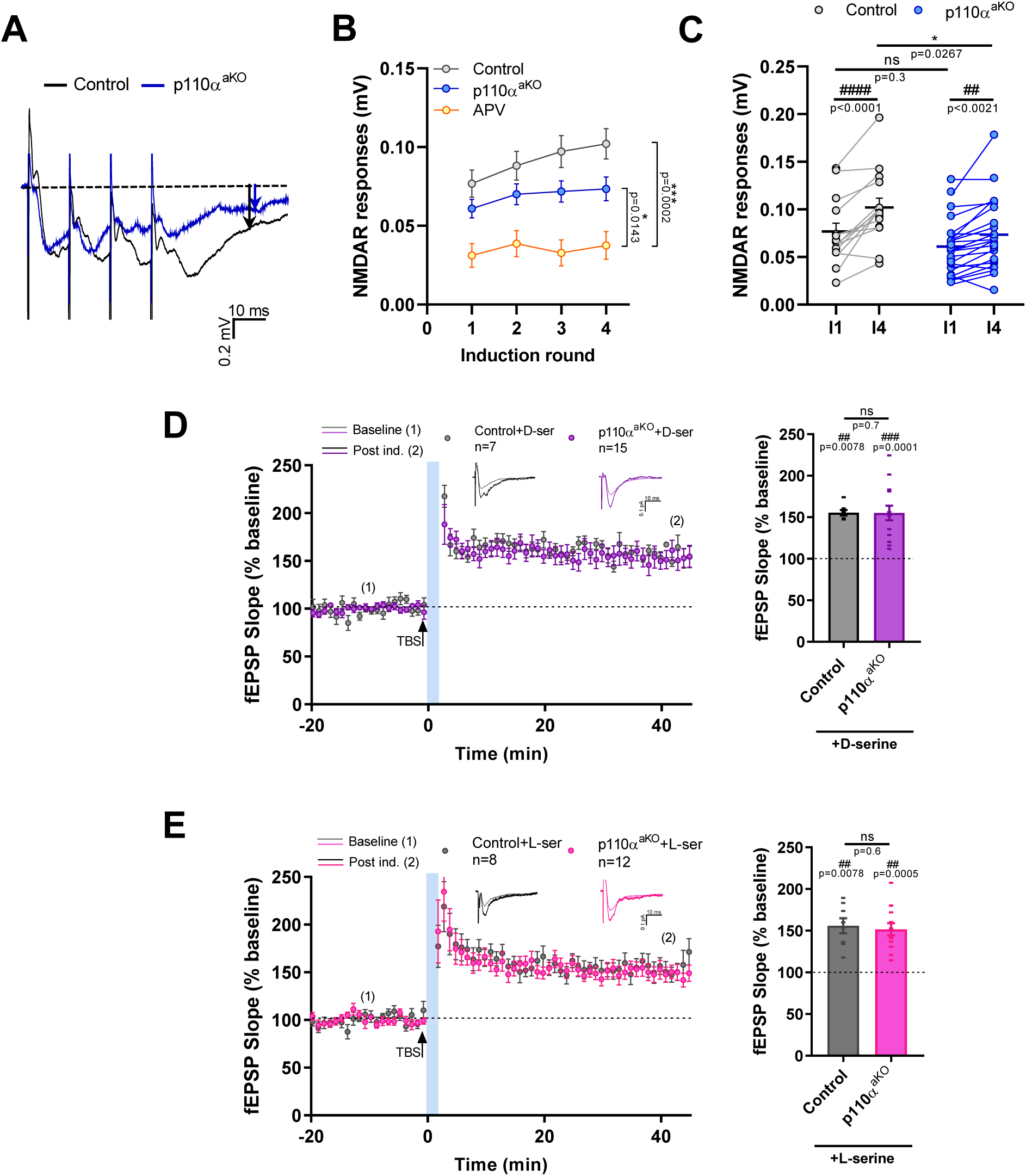
D- and L-serine treatments *in vitro* rescue the NMDAR-dependent LTP impairment in p110α astrocytic knock-out mice. **(A)** Average trace of 10 consecutive trains of 4 pulses for the last induction round for the control (black) and the p110α^aKO^ (blue) slices. The beginning of the dashed line represents the basal value and the arrows at 60 ms the fEPSP amplitude measured. The stimulation artifact for the 4 pulses can be seen before every peak of response. **(B)** NMDAR responses for the 4 induction rounds for p110α^aKO^, control and control APV treated slices. **(C)** NMDAR responses corresponding to the first and the fourth induction rounds in control and KO conditions. **(D, E)** Left, time course of NMDAR-dependent LTP induced by TBS with and without a treatment with 10 µM D-serine (D) or 50 µM L-serine (E). Right, bar plot representing the mean of the fEPSP slope for each slice on the last 5 min of recording. Representative traces for each condition are depicted in the upper part of the graph, the lighter color corresponding to the average trace of the baseline and the darker one to the post-induction. Bars represent mean±SEM and individual values for each condition are plotted as aligned dot-plots. Two-way ANOVA for matched samples and Bonferroni post-hoc were used in B and C to assess statistical differences between the induction rounds of each condition (#) and between p110α^aKO^ and the control (*) and Mann-Whitney and Wilcoxon Signed Rank Tests was used in D and E (ns: not significant) (* p<0.05, *** p<0.001, ## p<0.01, ### p<0.001, #### p<0.0001, ns: not significant). Control n=15, p110α^aKO^ n=23, APV n=12 slices from 2-15 mice. n, number of slices per condition from n>4 mice per condition.

We then reasoned that this specific deficit may related to the availability of D-serine, which is the preferred co-agonist (in addition to glutamate) for the activation of synaptic NMDARs during LTP^54^. In addition, astrocytes are good candidates to control D-serine availability, either from the conversion of L-serine into D-serine by the serine racemase (although this point is still controversial; see^55–58,60^) or from the biosynthesis of L-serine, which in the brain specifically occurs in glial cells^22,23,61,62^. To test whether the LTP impairment in the absence of astrocytic p110α was due to a deficit in D-serine availability, we carried out LTP experiments in the presence of 10 µM D-serine in the aCSF during the recordings. Under these conditions, the slices from p110α^aKO^ mice had normal LTP, comparable to controls also in the presence of D-serine (Fig. 4D). Then, to address whether this recovery of LTP was reflecting a specific deficit in D-serine production or a deficient biosynthesis of its precursor, L-serine, we tried these same rescue experiments with 50 µM L-serine in the aCSF during the recordings. Remarkably, the slices from p110α^aKO^ mice in presence of L-serine showed levels of LTP comparable to those seen in the treated controls (Fig. 4E). To note, the extent of LTP expression in control slices was similar with or without D- or L-serine (compare Fig. 3A with Fig. 4D, E), arguing against a general enhancing effect of these metabolites on LTP. Also, neither D-serine nor L-serine altered basal synaptic transmission in p110α^aKO^ animals as compared to controls (Suppl. Fig. 2A, B). Therefore, these combined results strongly suggest that the impairment in NMDAR activation during LTP induction in the absence of astrocytic p110α is due to a deficient biosynthesis of L-serine in astrocytes.

### The lack of p110α decreases glycolytic flux and increases mitochondrial respiration in astrocytes

As mentioned above, it is well established that the PI3K pathway constitutes a central hub for the regulation of glycolysis^33,37,41,42^. In addition, one of the major routes for L-serine biosynthesis is the phosphorylated pathway, which starts from the glycolytic intermediate 3-phosphoglycerate^22,59^. Therefore, it is reasonable to hypothesize that the absence of p110α in astrocytes leads to a reduced L-serine biosynthesis because of a lower glycolytic flux. To test this hypothesis, we evaluated glycolytic function at two points: glucose uptake and lactate production.

First, we assessed glucose uptake from the incorporation of the glucose analog 2-deoxyglucose in pure astrocytic cultures from p110α^flox/flox^ mice infected with an adenovirus expressing Cre, an empty vector (EV) or an uninfected condition (Suppl. Fig. 3A, B; see Materials and Methods). As shown in Fig. 5A, p110α^aKO^ astrocytes displayed a significant reduction of glucose uptake, compared to both EV-infected and uninfected astrocytes. Then, we analyzed lactate secretion as a readout of glycolytic flux by measuring the lactate concentration in the culture media. Consistent with the reduction in glucose uptake, we observed a significant decrease of lactate secretion to the media by the p110α^aKO^ astrocytes (Fig. 5B). These combined results suggest that p110α is an important driver of glycolysis in the astrocyte, starting from the uptake of glucose and producing a lower lactate yield.

**Figure 5.**
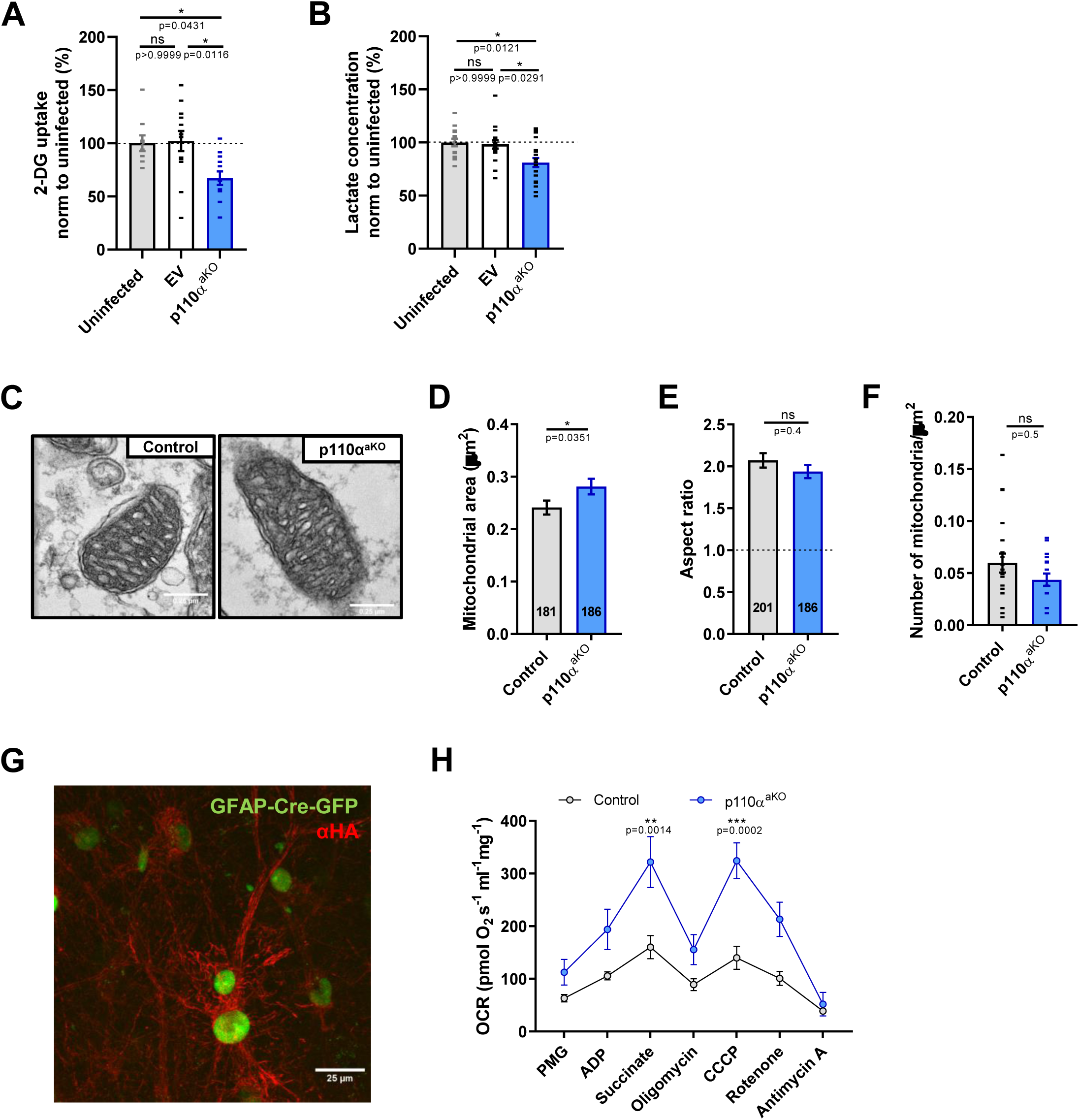
The lack of p110α in astrocytes decreases glucose uptake and lactate production in the astrocyte, while increasing mitochondrial size and function. **(A)** 2-DG uptake from astrocyte cultures infected with the adenovirus expressing Cre, with the empty vector (EV) or uninfected. All results are normalized to the 2-DG uptake in uninfected astrocytes. Uninfected n=9, empty vector n= 13 and p110α^aKO^ n=12 samples. **(B)** Extracellular lactate concentration levels from astrocytic cultures media normalized to the lactate concentration in the uninfected condition. Uninfected n=15, empty vector n= 18 and p110α^aKO^ n=20 samples. (A, B) bars represent mean±SEM and individual values from each sample are represented as a dot-plot overlay. Statistical differences were assessed by Kruskal-Wallis and Dunn’s multiple comparisons tests. **(C)** Electron microscopy representative images of astrocytic mitochondria from control (left) and p110α^aKO^ (right) mice from the *stratum radiatum* of acute hippocampal slices. Scale bar: 0.25 µm. **(D)** Mitochondrial area. Control n=181, p110α^aKO^ n=186 mitochondria). **(E)** The aspect ratio corresponds to the relation between the major and the minor axis of the mitochondria. Control n=201, p110α^aKO^ n=186 mitochondria. **(F)** Number of mitochondria per area. Control n=23, p110α^aKO^ n=18 sections. (D-F) n=3 mice per condition. Bars represent mean±SEM and individual values from each sample are represented as a dot-plot overlay. Statistical differences were assessed by Mann-Whitney test. **(G)** Merged representative image of the coinfection with the Cre-GFP AAV under the GFAP promoter and the AAV-mitoTag in the *stratum radiatum* of mouse organotypic slices. Cre-positive nuclei are shown in green and anti-HA labelled mitochondria in red. Scale bar: 25 µm. **(H)** OCR values normalized to protein levels from control and p110α^aKO^. Control n=4, p110α^aKO^ n=5 mice. Statistical differences were assessed by two-way ANOVA matched-values and Bonferroni post-hoc tests (* p<0.05, ** p<0.01, *** p<0.001). PMG, pyruvate, malate, glutamate; CCCP, carbonyl cyanide 3-chloropheny.

To further test the role of p110α in the regulation of metabolism in astrocytes, we decided to check the morphology and function of the mitochondria in p110α^aKO^ animals. For that purpose, we used the p110α^flox/flox^ mice injected with the AAV5-GFAP-Cre-GFP and their genotype-paired controls. After one month of expression, we prepared hippocampal sections and processed them for electron microscopy. We specifically imaged the CA1 *stratum radiatum* and identified astrocytes from their clearer cytoplasm and glycogen granules. As shown in Fig. 5C, D, astrocytes from mice lacking p110α had slightly bigger mitochondria than controls, without changes in their shape (aspect ratio values; Fig. 5E) or density (Fig. 5F). Also, mitochondria cristae were slightly longer and thinner in p110α^aKO^ mitochondria, without differences in number (Suppl. Fig. 4).

To evaluate mitochondrial function in the absence of astrocytic p110α, we measured the oxygen consumption rate from isolated mitochondria of astrocytes. For these experiments we took advantage of the Cre-dependent mitoTag adeno-associated viral vector (AAV-mitoTag) that expresses a recombinant TOMM20 protein fused to three hemagglutinin A (HA) in a Cre-dependent manner^47^. We coinfected p110α^flox/flox^ mice with AAV5-GFAP-Cre-GFP and AAV-mitoTag. In this manner, the Cre recombinase will drive the deletion of p110α from astrocytes and will tag their mitochondria through the HA-TOMM20 expression (see Fig. 5G for a representative image of coinfected astrocytes expressing GFP-Cre –green channel- and showing HA-labeled mitochondria –red channel). For these experiments, the controls were wild type mice coinfected with the same AAVs, so that the Cre expression will drive astrocytic mitochondrial tagging, but will not induce the p110α knock-out. The mitochondrial isolation was performed using whole hippocampal extracts incubated with anti-HA coated magnetic microbeads (see Materials and Methods). Using these preparations, we observed that astrocytic mitochondria from p110α^aKO^ mice had a marked enhancement of oxygen consumption rate compared to the wild type controls. This enhancement was statistically significant after the addition of the complex II substrate succinate and for maximal respiration driven by the mitochondrial uncoupler CCCP (Fig. 5H).

All together, these results point to an altered metabolism of astrocytes in the absence of p110α, which results in a defective glycolytic flux and an increase in mitochondrial respiration (perhaps as a compensatory mechanism for the glycolytic deficiency).

### L-serine supplement in vivo rescues the behavioral deficits in p110α^aKO^ mice

After strengthening the notion of a metabolic deficit in p110α^aKO^ mice and having observed that L-serine treatment *in vitro* was enough to rescue their LTP impairment, we decided to test whether the cognitive deficits in these animals could be rescued *in vivo*. To this end, we supplemented the mouse diet with 5 g/L of L-serine dissolved in the drinking water for three weeks, with the treatment starting after two weeks of infection and lasting until all behavioral assays were completed (Fig. 6A). After the behavioral assays, animals were sacrificed and D-serine was measured from total hippocampal extracts. Indeed, the treatment with L-serine significantly increased the levels of D-serine in p110α^aKO^ treated animals, which were now similar to those of control animals (Fig. 6B). These results indicate that the *in vivo* L-serine treatment is effective in increasing D-serine concentration in p110α^aKO^ animals.

**Figure 6.**
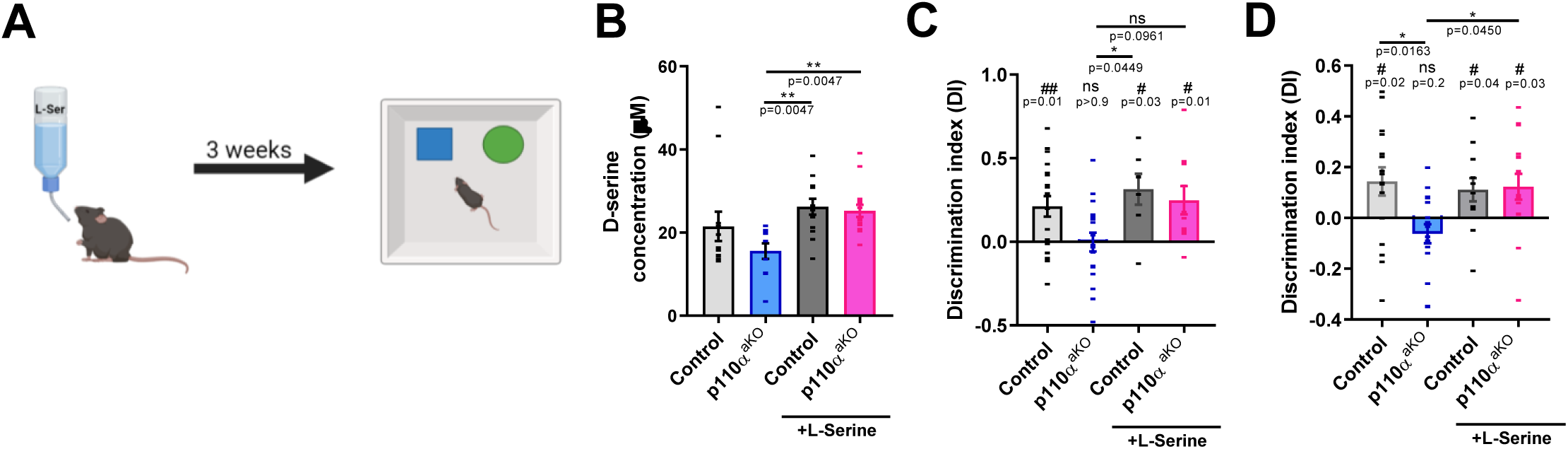
Spatial and object recognition memory in p110α^aKO^ mice are rescued with a supplement of L-serine in the drinking water. **(A)** Scheme depicting the 3 weeks L-serine treatment in the drinking water (5 g/L). **(B)** D-serine levels in total hippocampal extracts. Control n=12, p110α^aKO^ n=10, control+L-serine n=13, p110α^aKO^+L-serine n=16 mice. **(C, D)** Discrimination index for the novel location (C) or for the novel object recognition (D). Control n=17-19, p110α^aKO^ n=17-18, Control+L-ser n=7-12, p110α^aKO^+L-ser n=10-16 mice. The untreated controls and untreated p110α^aKO^ mice DIs are replotted from Fig. 2D, E, as reference. Individual values for each mouse and condition are represented along with the mean±SEM as aligned dot-plots. (C, D) To assess whether the DIs were different from zero we performed the Wilcoxon Signed Rank Test (# p<0.05, ## p<0.001). One-way ANOVA was used to assess statistical differences between the DIs and the D-serine concentration levels of p110α^aKO^, p110α^aKO^+L-ser, control and control+L-ser conditions (** p<0.01, * p<0.05, ns: not significant).

We then tested the behavioral assays in which we had found p110α^aKO^ animals to be deficient, namely NOR and NOL tests (Fig. 2D, E). For the NOL test, the performance of treated p110α^aKO^ animals was significantly higher than that of untreated animals, and was now similar to the discrimination of control animals with or without treatment (Fig. 7C). Similarly, in the case of the NOR test, the p110α^aKO^ mice that had received the L-serine supplement discriminated significantly more the new object over the familiar one, with a discrimination index similar to that of treated or untreated control animals (Fig. 7D).

These results indicate that a supplement of L-serine in the drinking water is enough to recover the cognitive deficits of mice lacking the p110α catalytic subunit of the PI3K in astrocytes. Also, they reinforce the link between LTP processes and learning, as well as the role of L-serine from the astrocyte for the correct functioning of the neuronal circuits, linking astrocytic metabolism to their modulation of synaptic activity.

## DISCUSSION

Astrocytes play a pivotal role not only through gliotransmission and modulation of synaptic responses, but also in energetic and metabolic regulation. This work sheds light on this matter, unraveling the distinct contributions of astrocytic PI3K signaling, and specifically the p110α isoform, to synaptic plasticity and cognitive function, highlighting its connection with metabolic regulation. In addition to the knowledge on basic mechanisms of synaptic plasticity, we believe this study provides further insight on brain disorders related to the PI3K pathway^31,63^ and/or D- or L-serine deficiencies^27,64^.

Perhaps one of the major contributions of this study is the dissection of specific mechanisms by which complex signaling pathways, with pleiotropic outcomes, may impinge on highly specialized cellular functions, such as synaptic plasticity and neuronal communication. The case of PI3K signaling is particularly paradigmatic. This pathway is long-known to be important for LTP^65–68^. Within neurons, its role is related to AMPAR synaptic delivery and anchoring^29,32,67^. However, we now show that this signaling molecule also contributes to LTP “at a distance” from the astrocyte, based on a completely different cellular function: regulation of glycolytic flux with the concomitant modulation of L-serine biosynthesis and D-serine availability for NMDAR activation during LTP. Of course, these distinct roles at neurons and astrocytes are relevant for pharmacological studies, where PI3K may be blocked broadly across the tissue (for instance, in some early studies^65–67^), but also when considering genetic mutations associated to human disease (for example, related to intellectual disabilities^31^), where the same mutation in different cell types (neurons, astrocytes) may contribute in distinct manners to the pathology.

In this sense, it is interesting that such a broad alteration in astrocytes (reduced glycolytic flux) may produce a precise phenotype in neurons: impaired long-term potentiation. Indeed, we have observed that other parameters of synaptic communication (glutamate release, basal synaptic transmission –mediated by AMPARs and NMDARs, or long-term depression) were not altered. This is likely reflecting the strong dependence on D-serine for LTP^54,55,58^. In fact, it has been shown that D-serine is not required for LTD and may even inhibit it^69^. Nevertheless, this specific effect on LTP does have additional effects on neurons. Thus, we also observed that deletion of astrocytic p110α produced a reduction in neuronal spine density with a concomitant increase in their length. These effects may reflect the presence of more immature spines, perhaps as a consequence to the failure to be potentiated and stabilized^70,71^. In turn, the reduced number of spines and their immature nature may affect their capacity to be potentiated and would ultimately impact in learning and memory. Importantly, this decrease in dendritic spine number and their lack of maturity could also be a consequence of the decreased lactate production in the p110α^aKO^ astrocytes, as it has been shown that lactate is important for LTP and spine dynamics^25,72^.

On the other hand, the specific alteration responsible for the reduced glycolysis in astrocytes lacking p110α remains to be determined. The PI3K/AKT pathway has been shown to regulate glycolytic flux in other tissues^33,41,73,74^. Interestingly, the knock-out for *Akt2*, the AKT isoform present in astrocytes^75^, is a common model for a type 2 diabetes-like syndrome, which shows an impairment in LTP and spatial memory^76,77^. Moreover, insulin resistance, a hallmark of type 2 diabetes, is linked to memory deficits and synaptic plasticity dysregulation in neurodegenerative diseases, aging or Alzheimer’s disease^78–80^, and astrocytic insulin receptor knock-out mice showed blunted responses to insulin and reduced brain glucose uptake^81,82^. This background suggests that the p110α^aKO^ might exhibit a diabetes-related phenotype, leading to a glycolytic deficiency, coursing with a decrease in glucose uptake and decrease in lactate production, which finally causes an impairment in long-term potentiation and memory.

It is also notable that astrocytes with reduced glycolytic flux in the absence of p110α displayed enhanced mitochondrial respiration. While astrocytes are considered mainly glycolytic cells, the role of mitochondria in astrocytes should not be overlooked, taking into account the differentially regulated enzymes and the distinct super-assembly of electron transport chain complexes between neurons and astrocytes^83,84^. In fact, previous p110α knock-out studies in muscle and adipose tissue have shown enlarged mitochondria with enhanced oxidative capacity^85,86^, in line with our observations in astrocytes. These results suggest a possible functional reorganization and adaptation of these knock-out astrocytes towards a more oxidative profile to sustain their own metabolic demands, possibly via fatty acid oxidation^83,87^.

Finally, the fact that the cognitive defect of p110α^aKO^ mice can be rescued *in vivo* by L-serine supplementation opens the possibility of dietary interventions with therapeutic potential. Indeed, L-serine is used as a routine food additive and dietary supplement, and it is well tolerated even at high doses for therapeutic applications^64,88^.

In summary, the results presented in this work highlight the dual role for p110α in orchestrating synaptic plasticity and metabolism in astrocytes in the hippocampus, with functional consequences for cognitive performance. Also, the application of an *in vivo* treatment to rescue the behavioral deficits paves the way for possible therapeutic strategies concerning neurological traits based on NMDAR hypofunction.

## ACKNOWLEDGMENTS

We thank the members of the Esteban laboratory for critical reading of the manuscript and the personnel at the fluorescence microscopy facility (SMOA), the electron microscopy facility and the animal house of the CBM for expert technical assistance. We also thank Cristina García-Cáceres and her team for technical advice for the glucose uptake assay protocol. We thank Susana Cadenas too, for performing the first Oroboros trials with total hippocampal homogenates.

## AUTHOR CONTRIBUTIONS

Conceptualization and experimental design: A.F.-R., M.I.C. and J.A.E. Most experimental work and data analysis: A.F.-R. Mitotag stereotaxic injections and oxygen consumption rate experiments: A.F.-R., A.E.-P and G.M. Assistance with some of the stereotaxic injections and behavioral assays: C.S.-C. Assistance with some of the biochemical analyses: E.L.-M., C.G.-V. and E.M. Assistance with astrocytic cultures: C.G.-V. Glucose uptake assay: C.G.-V. and S.G.-E. Neuronal morphology analysis: I.M. Extracellular lactate measurements: S.G.-E and C.B. Mouse genotyping: S.G.-E. Provision of p110α^flox/flox^: M.G. Provision of mitotag: A.Q. Writing original draft and editing: A.F.-R., M.I.C. and J.A.E.

