## Supplementary figures and images for "Astrocytic PI3Kα controls synaptic plasticity and cognitive function via serine metabolism"

### Supplementary figures

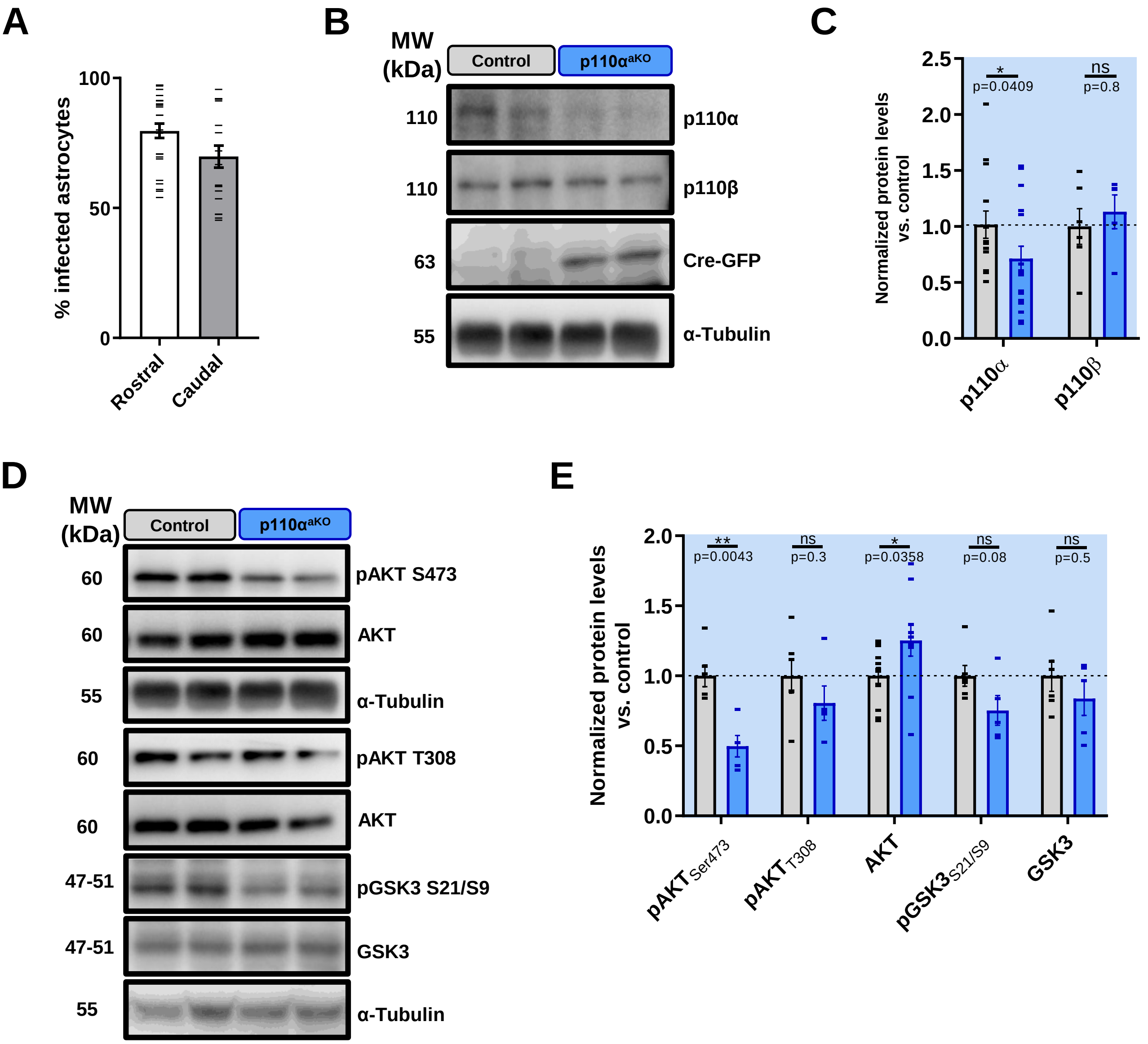


#### SUPPL. FIGURE 1

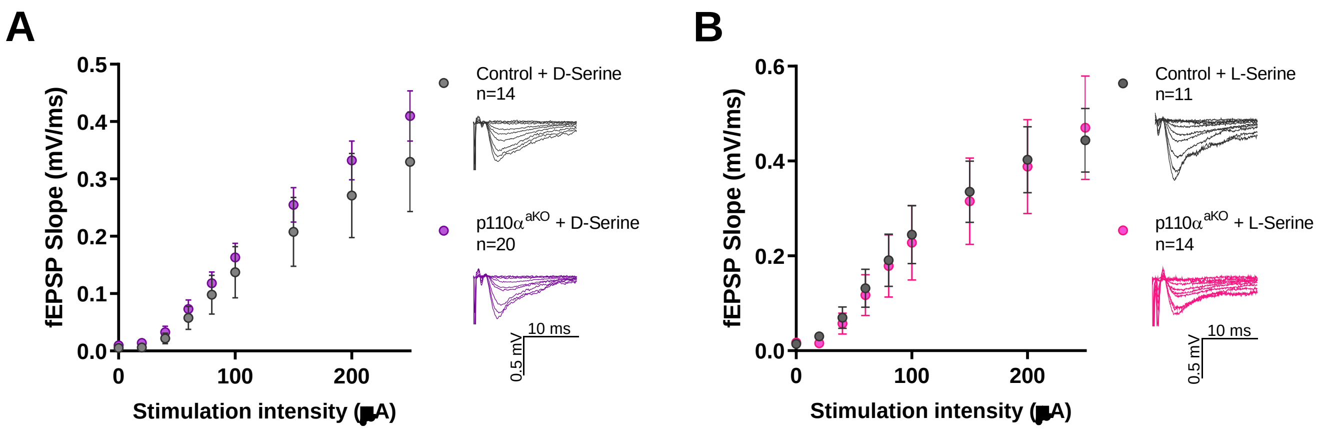


#### SUPPL. FIGURE 2

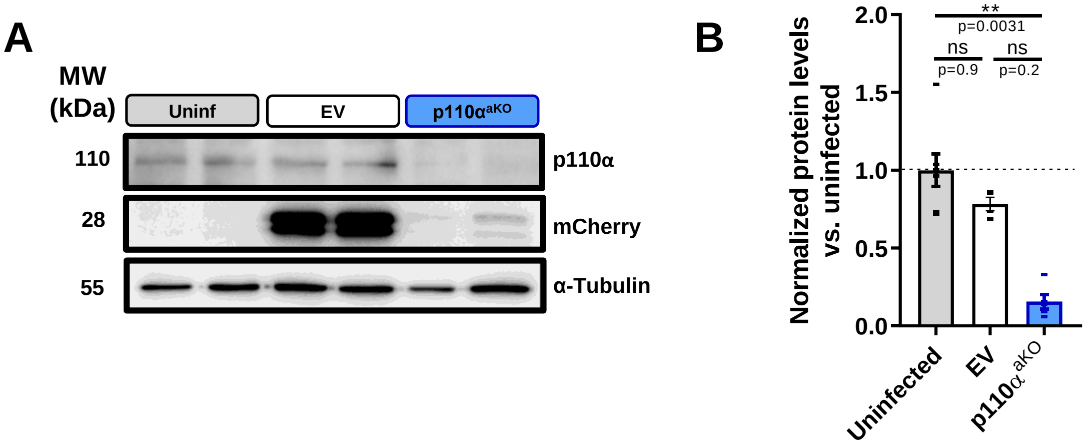


#### SUPPL. FIGURE 3

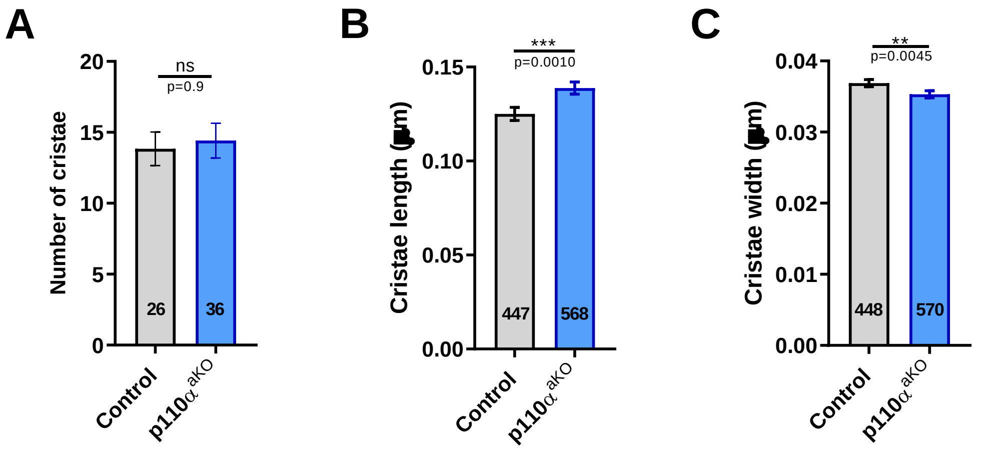


#### SUPPL. FIGURE 4
