## Supplementary text for "Astrocytic PI3Kα controls synaptic plasticity and cognitive function via serine metabolism"

### SUPPLEMENTARY INFORMATION

**Supplementary Figure 1. Biochemical analysis of p110α$^{aKO}$ hippocampal slices. (A)** Quantification of the fraction of infected astrocytes (green labeled nuclei) over the total number of astrocytes (GFAP staining) on the SR layer. The percentage of infected astrocytes was quantified along the rostro-caudal axis of the hippocampus to assess the targeting efficiency of the technique. n=26 rostral and 19 caudal slices from 19 different animals. **(B, D)** Representative western blots of whole hippocampal extracts of *in vivo* infected animals with the AAV driving the expression of Cre-GFP in astrocytes. Membranes were probed for p110α, p110β, GFP (infection level control) (B) and for downstream targets of the PI3K pathway such as $pAKT_{S473}$, $pAKT_{T308}$, total AKT, $pGSK3_{S21/S9}$ and total GSK3 (D). α-Tubulin was used as a loading control. **(C)** Quantification of the protein levels on p110α$^{aKO}$ mice and their saline-injected controls. Protein levels were normalized to α-Tubulin and then to the mean protein levels from the control samples. Control n=6-15 mice, p110α$^{aKO}$ n=6-19 mice. **(E)** Quantification of phosphorylation and total protein levels of p110α$^{aKO}$ vs. control. All the phosphorylated forms are normalized to their total and the total proteins are normalized to the loading control. All normalized protein levels are normalized to the mean levels from the control samples. Control n=5-6 mice, p110α$^{aKO}$ n=6 mice. (B, D, F) Bars represent mean±SEM. Individual values from each mouse are represented as a dot-plot overlay. Statistical differences were assessed by Mann-Whitney test. *p<0.05, **p<0.01, ns: not significant.

**Supplementary Figure 2. D- and L-serine treatments do not affect basal transmission in control and p110α$^{aKO}$ treated slices. (A, B)** Input-output curve of fEPSPs evoked at different stimulation intensities with 10 µM D-serine (A) or 50 µM L-serine (B) added to the electrophysiology bath. Representative traces from one experiment for each condition are shown in the right part of the graph. Scale bar: 0.5 mV, 10 ms. Statistical differences were assessed by two-way repeated-measures ANOVA with Bonferroni post-hoc test.

**Supplementary Figure 3. Conditional p110α knock-out in astrocytic cultures. (A)** Representative western blots of astrocyte culture extracts infected with the adenovirus expressing mCherry (empty vector) or mCherry-Cre (p110α$^{aKO}$). Membranes probed for p110α, mCherry and α-Tubulin as a loading control. **(B)** Quantification of the protein levels of p110α$^{aKO}$ vs. uninfected and EV vs. uninfected. Bars represent mean±SEM. Uninfected n=7, empty vector n=4 and p110α$^{aKO}$ n=5 samples. All proteins are normalized to their loading control and then to the mean levels from the control samples. Individual values from each sample are represented as a dot-plot overlay. Statistical differences were assessed by Kruskal Wallis and Dunn's multiple comparison tests (** $p<0.01$, ns: not significant).

**Supplementary Figure 4. Effect of p110α$^{aKO}$ in mitochondrial cristae. (A)** Number of cristae per mitochondria. Control n=26, p110α$^{aKO}$ n=36 mitochondria. **(B)** Cristae length. Control n=447 cristae from 28 mitochondria, p110α$^{aKO}$ n=568 cristae from 37 mitochondria. **(C)** Cristae width. Control n=448 cristae from 28 mitochondria, p110α$^{aKO}$ n=570 cristae from 37 mitochondria. n=3 mice per condition. Bars represent mean±SEM and individual values from each sample are represented as a dot-plot overlay. Statistical differences were assessed by Mann-Whitney test (** p<0.01, *** p<0.001, ns: not significant).
